# The effect of dietary ginseng polysaccharide supplementation on the immune responses involved in porcine milk-derived esRNAs

**DOI:** 10.1101/101592

**Authors:** Jiajie Sun, Liyuan Yao, Ting Chen, Qianyun Xi, Yongliang Zhang

## Abstract

Ginseng and its polysaccharides (GPS) have been well known as an immune modulator. This study was conducted to investigate the effects of dietary supplemental GPS on the immune responses involved in sow’s milk-derived exosomal shuttle RNAs (esRNAs) using RNA-Seq and miRNA-Seq. Of the 213 identified miRNA types, a total of 26 conserved miRNAs were differently expressed in response to GPS supplementation, including 10 up-regulated and 16 down-regulated miRNAs in GPS feeding group. In addition, exosomal transcriptome analysis identified 14,696 protein-coding genes in sow’s milk exosomes, and 283 genes with 204 and 79 candidates showing up and down-regulation were significantly responded to GPS supplementation. Integrated analysis of each differently expressed miRNA with significantly expressed genes further revealed the presence of 51 highly conserved miRNA-gene interactions that were annotated to be related to immunoregulatory functions. This work provided an important advance in the functional identification of dietary GPS supplementation and more fundamental information about how GPS promoted the immune response and healthy growth of the infant from mothers at molecular levels.

## Introduction

Ginseng (Panax ginseng C.A. Meyer), one of the most well-known oriental medicine for several thousand years, has been widely used with mysterious powers as a tonic, prophylactic and restorative agent, etc. (Sun 2011) The polysaccharide extracted from the medicinal ginseng root (mostly), stems and leaves was demonstrated to have many functions, including inhibition of tumors (Li *et al*. 2014), suppression of bacterial (Fukuyama *et al*. 2012) and viral (Kim *et al*. 2011) activity, anti-peroxidatic reactions (Luo et al. 2008), and innate (Shin *et al*. 2002) or acquired (Sumiyoshi *et al*. 2011) immune modulation. In recent years, there has been growing interests in the use of polysaccharides as new, alternative immunologic additives for agricultural animals. Chen et al. (Chen *et al*. 2009) indicated an increase in cellular and humoral immunities by modulating the production of antibodies, complements and cytokines in the Achyranthes bidentata polysaccharide supplemented weaned piglets, conferring an important protective role in the non-specific defense against infections. In our previous study, dietary GPS significantly increased immune enzyme activity and modified expression of immune genes in shrimp (Liu *et al*. 2011). However, to date the effects of dietary polysaccharide supplementation on lactating sows was deficient.

In general, breast milk supplied the chief nutrient source during the natural suckling period of the neonates, and milk liquid with immune-related composition also provided immunity to the infant and affected the maturation of the infant’s immune system (Admyre *et al*. 2007). Previous research has reported that exosomes were identified to present in the breast milk (Zhou *et al*. 2012), which were small membrane vesicles of endocytic origin that were released from the producing cell into the extracellular environment (Théry *et al*. 2002). And exosomes have been proposed to signal by binding to the recipient cell surface receptors or by internalisation with the cell membrane (Valadi *et al*. 2007), potentially donating substantial amounts of exosomal RNAs such as mRNAs, microRNAs (miRNAs), and other non-coding RNAs (ncRNAs) to other cells and subsequently affecting the protein production of a recipient cell (Sato-Kuwabara *et al*. 2005). In addition, previous research has revealed the ability of human breast milk exosomes to potentially influence the immune system of the infant (Admyre *et al*. 2007).

Overall, we therefore hypothesized that dietary supplementation with GPS influenced the composition of sow milk, especially for immune-related esRNAs, and further enhanced the immune responses in suckling piglets. This hypothesis was performed by miRNA and RNA sequencing to analyze the porcine breast milk exosomes in response to GPS supplementation.

## Materials and Methods

### Animals and feeding

The GPS were extracted from the roots of P. ginseng according to the protocols described in our previous research (Liu *et al*. 2011). A total of 20 Large White sows at 90 days of gestation were acquired from WENS breeding pig farm (Qingyuan, Guangdong, China) and allocated into two dietary treatment groups on the basis of age, body weight and parities. Control group (Con) was fed the basal diet, and the diet of treatment group was added with 400 mg/kg GPS until 14th day after delivery. Porcine milk samples were collected at day 1, 3, 7 and 14 after parturition, and kept at -80°C until use.

### Milk exosomes collection and sequencing analysis

Preparation of exosomes from porcine milk was operated as our previous descriptions (Chen *et al*. 2014). Total RNAs were extracted from pelleted exosomes using TRIzol^®^ Reagent (Life Technologies, Carlsbad, CA, USA) by the manufacturer’s protocol, and the quantity and purity were analysis using Bioanalyzer 2100 and RNA 6000 Nano Labchip Kit (Agilent, CA, USA) with RNA Integrity Number (RIN) value ≥7.0. Prior to constructing Illumina-indexed libraries, the total RNAs from four different time points were pooled equally in each treatment group. Subsequently, indexed miRNAome and transcriptome sequencing libraries were constructed with Illumina TruSeq Small RNA Library Preparation Kits and TruSeq RNA Library Preparation Kits (Illumina, San Diego, CA) respectively, which included size selection of the final library amplicons. Finally, the libraries were sequenced using Illumina HiSeq^TM^ 2000, and the raw reads were demultiplexed and the indexed adapter sequences were trimmed using the CASAVA v1.8.2 software.

### miRNA analysis

Firstly, the raw reads of control and GPS supplemented group were processed with ACGT101-miR program (LC Sciences, Houston, Texas, USA) to trim 3’ adapter and discard reads with poly N and reads shorter than 17 bp. Then, using BLAST search all clean reads were aligned to porcine miRNA mature and precursor sequences published in miRBase v21.0 (http://www.mirbase.org/) to identified conserved miRNAs. The read count of each identified miRNA was firstly normalized to reads per million, and the R v3.0.2 Bioconductor package EDGER v2.4.6 (Robinson *et al*. 2010) was applied to identify differentially expressed (DE) miRNAs with p value < 0.05 and fold change ≥ 2 between different groups.

### Align the RNA-seq reads to genome and assemble expressed transcripts

Raw reads of fastq format were firstly processed by removing the low quality reads and the reads that contain adapter or ploy-N. Then, the clean reads from each library were aligned to the Sscrofa10.2 reference genome (http://hgdownload.soe.ucsc.edu/goldenPath/susScr3/bigZips/susScr3.fa.gz) downloaded from the the University of California Santa Cruz (UCSC) website with TopHat v2.0.12 (Trapnell *et al*. 2009). The mapped reads of each group were assembled by Cufflinks 2.2.1 (Trapnell *et al*. 2012), and using Cuffmerge two assemblies were merged to create a single transcriptome annotation with porcine Ensembl’s genes generated by UCSC table browser for subsequent protein-coding gene analysis. The gene expression level was analyzed by using RPKM method (Reads per kilobase transcriptome per million mapped reads) (Mortazavi *et al*. 2008), and only genes with False Discovery Rate (FDR) value < 0.05 that calculated by Cuffdiff program and |log2 fold change|≥1 were considered as differently expressed candidates.

### miRNA target prediction and network construction

Putative targets of DE miRNAs were evaluated using miRanda (Betel *et al*. 2010), and only alignments with energies ≤ -20.0 kcal/mol and no mismatch in the seed region (positions 2–8 in the 5′end) were used for further analysis. The construction of interaction networks included three steps: (□) DE mRNAs and DE protein-coding genes were first retained; (□) then mRNA-miRNA negative interactions were predicted by miRanda analysis; (□) potential interactions of targets-miRNA were established and visualized using Cytoscape V3.4 (http://cytoscape.org/).

## Results and Discussion

### Sequence analysis of milk exosomal miRNAs

Solexa sequencing provided a total of 13,256,489 and 11,349,485 raw reads of 51 nt from the Con and GPS libraries, respectively. After removing 3′ adapter null or 5′ adapter contaminants, low quality reads, and reads contained poly N or smaller than 17 nt, a total of 12,256,073 and 10,654,756 clean reads of 17–31 nt were obtained (Table 1). Subsequently, clean reads were mapped to porcine miRNA mature and precursor sequences published in miRBase 21.0 by BLAST search to identify known miRNAs, and length variation at both 3′ and 5′ ends and one mismatch inside of the sequences were allowed in the alignment. In total, we identified 213 unique miRNA types aligned to 181 independent pre-miRNA loci with read number more than 10, and 188 known miRNAs were expressed in both Con and GPS groups (Table S1A). The 10 most highly expressed miRNAs in each group accounted for 89.64% ± 3.07% of the total counts of all unique miRNAs, and four of these miRNAs (ssc-miR-148a-3p, ssc-miR-30a-5p, ssc-miR-21 and ssc-miR-26a) were found in common across two tested groups, indicating that the majority of miRNAs were expressed from very few loci that maybe play very important roles in milk. Recently, similar miRNAs were identified by sequencing of porcine (Chen *et al*. 2014), bovine (Sun *et al*. 2015) and human (Chen *et al*. 2010) milk exosomes, and miR-148a-3p and miR-30a were also represented as the top 10 miRNAs. Of the identified miRNAs, the most abundant was miR-148a-3p, which represented 56.02% ± 5.88% reads across two libraries. The predominance of miR-148a was consistent with its well-established function as a nutritional biomarker corresponding to the protein content of various bovine-derived milk products (Chen *et al*. 2010). Interestingly, feeding studies that have shown exogenous plant (Zhang *et al*. 2012) or milk (Baier *et al*. 2014) miRNAs can be found in the sera and tissues and influence regulation of target genes in recipient animals, suggesting that the enrichment of specific miRNAs derived from milk in our study may also influence the growth and development of neonates. Specifically, miR-148a influenced dendritic cell activation and maturation by down-regulating calcium/calmodulin-dependent protein kinase IIa (CaMKIIa) expression, and finally associated with anti-inflammatory responses (Turner *et al*. 2011). MiR-30a attenuated immunosuppressive functions of IL-1β-elicited mesenchymal stem cells via targeting transforming growth factor-β-activated kinase 1 binding protein 3 (TAB3) (Hu *et al*. 2015). Also widely studied for possible involvement in the pathogenesis of multiple autoimmune and chronic inflammatory disorders, miR-21 expression was specifically elevated in Th17 cells, and T cell-intrinsic expression of miR-21 was important for effective Th17 differentiation (Murugaiyan *et al*. 2015). In addition, cellular miR-26a suppressed replication of porcine reproductive and respiratory syndrome virus by activating innate antiviral immunity (Jia *et al*. 2015). Even though there were compelling evidences that the most prevalent miRNAs in our study could potentially exert an influence on immune response, the specific functional roles of these miRNAs need further detailed investigations to obtain a thorough understanding of the specific targets and mechanistic effects on consumption of miRNA-loaded porcine milk exosomes by a recipient animal.

**Table 1.**
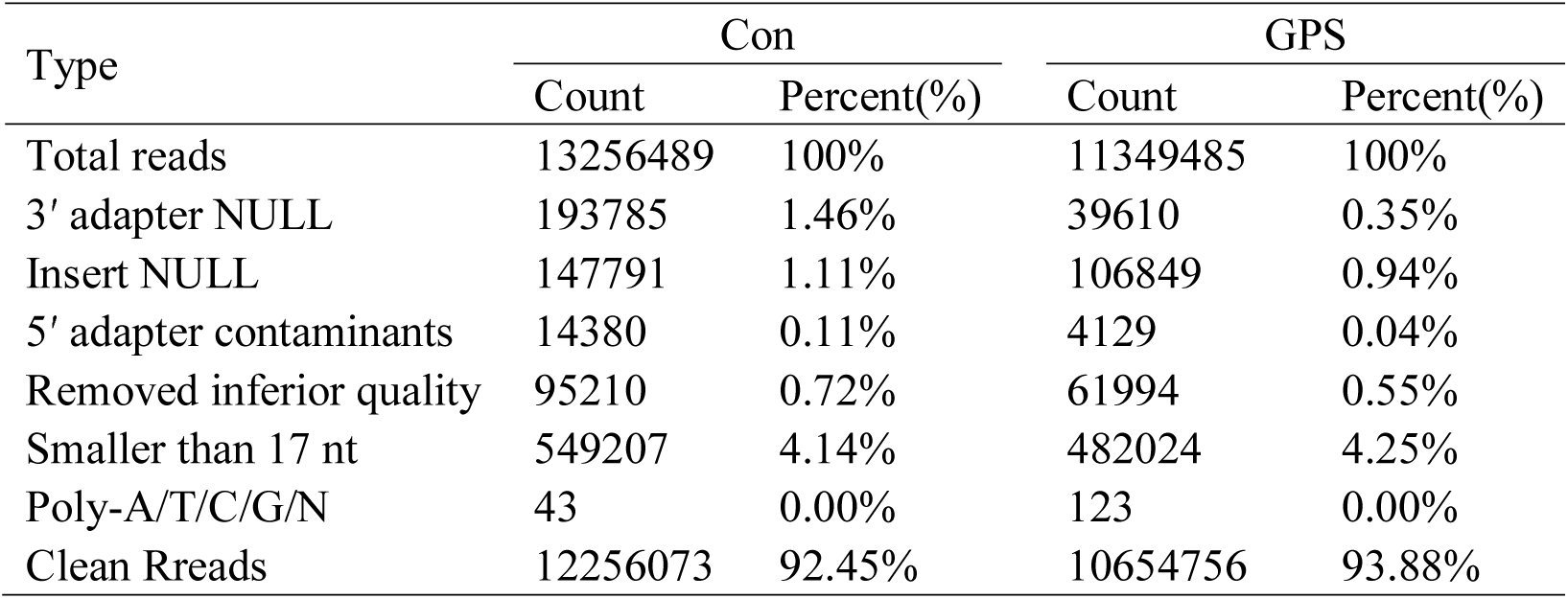
Summary of small RNA sequencing date

### Expression analysis on porcine milk exosomal transcriptome

The single-end RNA reads generated from sequencing of the exosomal libraries were trimmed to remove adapter sequences and then filtered. After filtering procedures, a total of 77,106,888 and 82,953,484 high-quality reads were considered for further analysis, resulted in ~6.46 and ~6.95 gigabases (Gb) from Con and GPS libraries, respectively. We aligned all these high-quality reads onto the porcine Sscrofa10.2 reference genome, and found that over 63.59% ± 0.25% of the reads were mapped to the genome, including 58.31% ± 0.23% of the mapped reads that were aligned uniquely in each library (Table S2A). Transcripts assembled with mapped reads using cufflinks revealed a total of 14,696 protein-coding genes across the two libraries, and Con specific units accounted for 95.30% of all identified genes, as well as 94.55% existed in the GPS library (Table S2B), consistent with previous study indicating that 19,320 mRNA transcripts were present in bovine milk exosomes (Izumi *et al*. 2015). In addition, the top 100 highly expressed genes in sequencing libraries accounted for 72.91% ± 0.96% of the total expression levels calculated by RPKM value, and a total of 98 genes were common to both groups. We then performed Gene Ontology (GO) enrichment analysis of these highly expressed genes using the DAVID bioinformatics resource, which employs a Fisher’s Exact Test with Benjamini–Hochberg correction. A total of 82 enriched GO categories were derived using a P-value cut-off of P < 0.05, including 56 Biological Process (BP), 19 Cell Component (CC) and 7 Molecular Function (MF) categories (Table S2C), respectively. These categories were mainly focused on translational elongation, ribosome biogenesis, rRNA processing and rRNA metabolic process, clearly indicating that the exosomal RNAs were enriched in rRNA-family species (Jenjaroenpun *et al*. 2013). Jenjaroenpun et al. found that cellular exosomes contained various classes of RNA moleculars with the major class represented by fragmented ribosomal RNA (rRNA), in particular 28S and 18S subunits. The observation in our and previous studies suggested that the majority of exosomal rRNAs were fragmented, and these results may explain why the 28S and 18S rRNA cannot be detected in exosomal total RNAs (Gu *et al*. 2012).

### miRNA and their target expression in response to GPS supplementation

After normalization, miRNAs that changed more than 2-fold and p value less than 0.05 were considered to be differently up or down regulated in response to GPS supplementation. In total, there were 10 miRNAs up-regulated in the milk exosomes of GPS group compared with control, as well as 16 down-regulated miRNAs (Table S1B). To identify the potential function of these DE miRNAs, target prediction was performed using miRanda software. Prediction analyses yielded a total of 3,525 unique genes potentially regulated by DE miRNAs. This resulted in 4,902 miRNA-target interactions; 3,271 of these were targeted by up-regulated miRNAs, and 1,631 were targeted by down-regulated miRNAs (Table S1C). The KEGG pathway analysis of the predicted targets by DAVID Bioinformatics Resources revealed 22 unique KEGG terms with P vaule < 0.05, which had been previously implicated in immune and disease. These pathways included the top 5 statistical enrichment, namely Pancreatic cancer, MAPK signaling pathway, B cell receptor signaling pathway, T cell receptor signaling pathway and Fc gamma R-mediated phagocytosis (Table S1D), suggesting the functions of GPS in immune enhancement and regulation.

Previous reports have documented the functional connection between RNA editing and miRNA-mediated post-transcriptional gene silencing (Bartel *et al*. 2009). To characterize the potential effects of RNA editing on miRNAs in porcine milk exosomes, we first profiled porcine milk exosomal transcriptome between Con and GPS groups. Of 14,696 identified protein-coding genes, a total of 283 exosomal genes were further considered to be differently expressed in response to GPS supplementation based on |log2 fold change|≥1, FDR < 0.05 and gene coverage≥60%, including 204 up-regulated and 79 down-regulated genes in GPS groups (Table S2D). Next, we aimed to compute all possible interactions of each DE miRNA with significantly expressed genes. Generally, an up-regulation of a target gene indicates a decrease activity of the corresponding miRNA; therefore, a miRNA-mRNA interaction pair means anti-regulation of a miRNA and a corresponding gene. In final, miRanda analysis revealed the presence of 51 highly conserved miRNA-mRNA interactions. Of these negative interactions, a total of 9 down-regulated genes appeared to be targeted by 8 up-regulated miRNAs in GPS group, and 17 up-regulated genes was targeted by 12 down-regulated miRNAs (Figure 1). By applying a cut-off criterion of FDR value < 0.05, GO enrichment analysis of these target candidates revealed a few important terms those were significantly enriched in the functions of immunity, such as neutrophil chemotaxis (CCL3L1, FCER1G, TGFβ2), phagocytosis (FCER1G, BIN2), cell chemotaxis (CXCL2, BIN2) and chemokine-mediated signaling pathway (CCL3L1, CXCL2) (Table S2E). In details, CCL3L1 dramatically influenced cell-mediated immunity (Dolan *et al*. 2007) and was targeted by ssc-miR-19a in our paper. Phagocytosis-related molecule FCER1G played a role in uptake of antigens bound to immunoglobulin E (Landi *et al*. 2011), and cytokine TGFβ2 acted an integral role in regulating immune responses (Rautava *et al*. 2011), these implying the biological functions of their anti-corresponding miR-22-3p and miR-374b-5p in porcine milk exosomes. In addition, kidney-specific miR-192 (Liang *et al*. 2007) significantly involved in immune-related gene expression (Wu *et al*. 2012), consistent with our results in negative regulation of BIN2, a membrane sculpting protein that influenced leucocyte podosomes, motility and phagocytosis (Sánchez-Barrena *et al*. 2012). And mast cell and macrophage chemokine CXCL2, a target gene of miR-30a-3p, controlled the early stage of neutrophil recruitment during tissue inflammation (De Filippo *et al*. 2013). Taken together, we found that the expression of several key miRNAs and genes in porcine milk-derived exosomes were induced or decreased under GPS supplementation in porcine basal diets, and these miRNA-mRNA interactions were reported to be significantly related to immunoregulatory functions. Meanwhile, previous research has showed that exosomal mRNAs or miRNAs, termed esRNAs, can be delivered to another cell and take effect on the new location (Valadi *et al*. 2007). Therefore, the findings suggested the functions of dietary GPS supplementation in enhancing the immune-related esRNAs expression in milk-derived exosomes, and these candidates may be further transferred to suckling piglets for immune system development and healthy growth.

**Figure 1.**
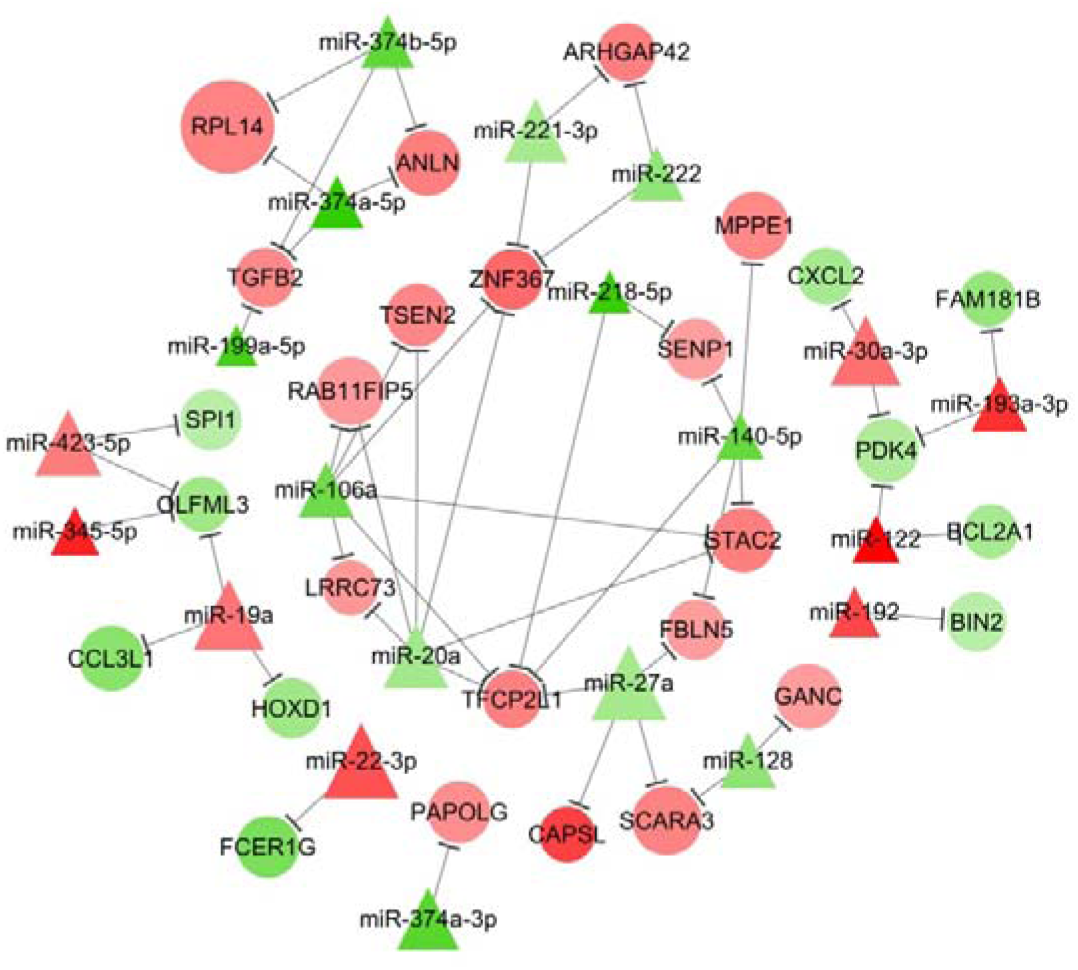
miRNA-mRNA correlation networks. Note: Circular nodes represented targets and triangular nodes represented miRNAs. Red nodes represented an up-regulation and green represented a down-regulation relative to match in GPS group. The size of each node represented for average expressed level of each gene between two groups, and the significant levels between different groups were associated with the color of node deepening.

In conclusion, we detected 26 miRNAs and 283 genes in porcine milk-derived exosomes that showed differential expression in response to GPS supplementation in sow’s basal diets. Integrated analysis and functional annotation revealed 51 highly conserved miRNA-mRNA interactions that were significantly related to immunoregulatory functions. This work provided an important advance in the functional identification of dietary GPS supplementation and more fundamental information about how GPS promoted the immune response and healthy growth of the infant at the molecular level.

## Acknowledgments

This work was supported by grants from the Natural Science Foundation of Guangdong province (2016A030310449, 2016A030313413), the Key Project of Transgenic Animal (2014ZX0800948B), the National Basic Research Program of China (973 Program, 2013CB127304), and Natural Science Foundation of China program (31272529, 31472163). None of the authors declare any conflicts of interests.

### Conflict of Interests

The authors have declared that no conflict of in-terest exists.

### Supporting information available

Known miRNAs identified in both Con and GPS groups were showed in Table S1A; differently expressed miRNAs between Con and GPS groups in response to GPS supplementation were showed in Table S1B; miRNA target gene prediction were showed in Table S1C; KEGG analysis of the target genes of DE miRNAs were showed in Table S1D. Summary of RNA-seq dates were showed in Table S2A; Protein-coding genes identified in both Con and GPS groups were showed in Table S2B; Gene Ontology (GO) enrichment analysis of highly expressed genes were showed in Table S2C; Differently expressed genes between Con and GPS groups in response to GPS supplementation were showed in Table S2D; GO enrichment analysis of target candidates presented in miRNA-mRNA interaction network were showed in Table S2E.

